# Computational prediction of drug response in short QT syndrome type 1 based on measurements of compound effect in stem cell-derived cardiomyocytes

**DOI:** 10.1101/2020.06.24.168690

**Authors:** Karoline Horgmo Jæger, Samuel Wall, Aslak Tveito

## Abstract

Short QT (SQT) syndrome is a genetic cardiac disorder characterized by an abbreviated QT interval of the patient’s electrocardiogram. The syndrome is associated with increased risk of arrhythmia and sudden cardiac death and can arise from a number of ion channel mutations. Cardiomyocytes derived from induced pluripotent stem cells generated from SQT patients (SQT hiPSC-CMs) provide promising platforms for testing pharmacological treatments directly in human cardiac cells exhibiting mutations specific for the syndrome. However, a difficulty is posed by the relative immaturity of hiPSC-CMs, with the possibility that drug effects observed in SQT hiPSC-CMs could be very different from the corresponding drug effect *in vivo*. In this paper, we apply a multistep computational procedure for translating measured drug effects from these cells to human QT response. This process first detects drug effects on individual ion channels based on measurements of SQT hiPSC-CMs and then uses these results to estimate the drug effects on ventricular action potentials and QT intervals of adult SQT patients. We find that the procedure is able to identify IC_50_ values in line with measured values for the four drugs quinidine, ivabradine, ajmaline and mexiletine. In addition, the predicted effect of quinidine on the adult QT interval is in good agreement with measured effects of quinidine for adult patients. Consequently, the computational procedure appears to be a useful tool for helping predicting adult drug responses from pure *in vitro* measurements of patient derived cell lines.

**Author summary:** A number of cardiac disorders originate from genetic mutations affecting the function of ion channels populating the membrane of cardiomyocytes. One example is short QT syndrome, associated with increased risk of arrhythmias and sudden cardiac death. Cardiomyocytes derived from human induced pluripotent stem cells (hiPSC-CMs) provide a promising platform for testing potential pharmacological treatments for such disorders, as human cardiomyocytes exhibiting specific mutations can be generated and exposed to drugs *in vitro*. However, the electrophysiological properties of hiPSC-CMs differ significantly from those of adult native cardiomyocytes. Therefore, drug effects observed for hiPSC-CMs could possibly be very different from corresponding drug effects for adult cells *in vivo*. In this study, we apply a computational framework for translating drug effects observed for hiPSC-CMs derived from a short QT patient to drug effects for adult short QT cardiomyocytes. For one of the considered drugs, the effect on adult QT intervals has been measured and these measurements turn out to be in good agreement with the response estimated by the computational procedure. Thus, the computational framework shows promise for being a useful tool for predicting adult drug responses from measurements of hiPSC-CMs, allowing earlier identification of compounds to accurately treat cardiac diseases.

## 1 Introduction

Short QT (SQT) syndrome is a cardiac channelopathy characterized by an abnormally short duration of the QT interval of the patient’s electrocardiogram (ECG) [1, 2, 3]. The syndrome is associated with increased risk of atrial fibrillation, ventricular arrhythmias and sudden cardiac death [4, 5] and was first described by Gussak et al. in 2000 [1]. The first identified SQT subtype, termed SQT1, results from an increase in the transmembrane potassium current, *I*_Kr_, caused by a mutation in the *I*_Kr_ encoding gene KCNH2 [6]. Later, several additional subtypes of SQT syndrome have been identified, originating from mutations in other genes, including gain of function alterations in genes encoding the potassium channels responsible for the *I*_Ks_ and *I*_K1_ currents and loss of function alterations in genes encoding calcium channels [2, 3].

Because of the lethal risks associated with the syndrome, there is an urgent need for effective pharmacological therapies. One modern approach is to identify compounds that can make the electrophysiological properties of cardiomyocytes (CMs) populated with SQT mutated channels more similar to the electrophysiological properties of CMs populated with wild type (WT), unmutated, channels (see e.g., [7, 8, 9, 10, 11]). For SQT1, which is characterized by an increased *I*_Kr_ current, drugs inhibiting the *I*_Kr_ current have been proposed as possible candidates (see e.g., [11, 12]). However, the effect of a drug on WT *I*_Kr_ channels can be very different from the effect on mutated channels. For example, the *I*_Kr_ blockers sotalol and ibutilide have been shown to be ineffective for SQT1 patients [12]. This may be explained by the fact that these drugs primarily affect the inactivated state of the *I*_Kr_ channels, and the SQT1 mutation impairs the inactivation of the channels [6, 13, 14]. On the other hand, quinidine, which affects both the open and inactivated states of *I*_Kr_ channels has proven to be more effective for SQT1 patients [15]. These examples demonstrate that investigations into drug effects on specific SQT mutations are needed in order to find appropriate pharmacological treatments.

A promising prospect in the quest for suitable drugs for SQT syndrome is the development of human induced pluripotent stem cells (hiPSCs) (see e.g., [16, 17, 18, 19]). These cells can be generated from individual patients and differentiated into a large number of different cell types, including CMs. Thus, cells can be generated from SQT patients, allowing for investigations of drug effects for CMs populated by ion channels affected by the patient specific mutations. Indeed, in [9, 11], several drugs have been tested on hiPSC-derived CMs (hiPSC-CMs) from an SQT1 patient, revealing possible promising drugs.

However, a difficulty with using hiPSC-CMs in drug testing applications is that the electrophysiological properties of hiPSC-CMs differ significantly from the electrophysiological properties of adult CMs (see e.g., [20, 21]). In general, hiPSC-CMs are recognized as electrophysiologically immature, with properties more similar to those of fetal CMs. These differences imply that the drug response observed for hiPSC-CMs might not be the same as the corresponding drug response for adult native CMs.

Mathematical modeling has been proposed as a possible tool to help translate the drug response of hiPSC-CMs to the drug response of adult CMs (see e.g., [22, 23, 24, 25]). The development of mathematical models of the dynamics underlying human cardiac action potentials is an active field of research, and a large number of models have been developed, including models for adult undiseased ventricular CMs [26, 27], atrial CMs [28, 29], hiPSC-CMs [30, 31] and CMs affected by mutations [32, 33]. In these models, changes in the membrane potential are represented by individual transmembrane ionic currents (see e.g., (1) below), and the models can therefore be useful for investigating how drug effects on individual ion channels affect the composite dynamics of the full action potential. Furthermore, the action potential models can be combined with spatial models of electric conduction (see e.g., [34, 35, 36, 37, 38]) to provide insight into drug effects on cardiac conduction properties and mechanisms of arrhythmia.

In previous studies, we have applied a procedure based on mathematical action potential modeling to estimate drug effects on individual ion channels from measurements of hiPSC-CMs and predict adult CM drug responses [22, 23]. This procedure is based on the assumption that the function of individual proteins, e.g. ion channels, is the same for hiPSC-CMs and adult CMs, and that only the density of the proteins differs between hiPSC-CMs and adult CMs (see Figure 2). From this assumption, it follows that the effect of a drug on an individual ion channel is the same for hiPSC-CMs and adult CMs. Assuming that we have correctly identified the drug effect on individual channels in the hiPSC-CM case, we can therefore directly translate this effect to the adult case by inserting the inferred mechanisms into a model for adult cells.

**Figure 1:**
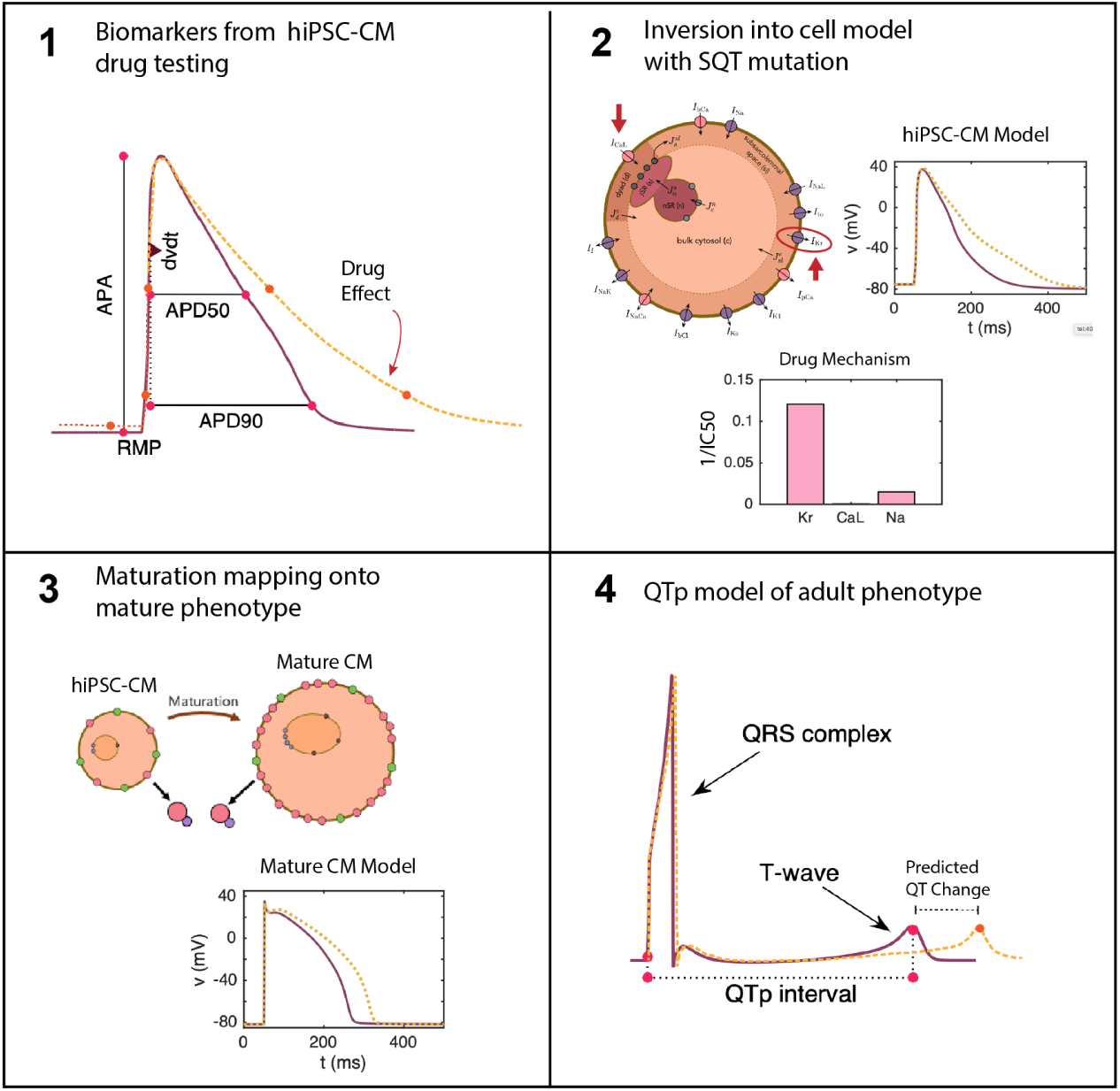
Illustration of the computational pipeline. 1) Biomarkers from the cardiac AP are taken from hiPSC-CMs under drug testing. 2) These biomarkers from dose escalation studies are inverted into an SQT1 model of the AP of hiPSC-CMs. Inversion into a matched model provides determination of drug effects on specific channels [23]. 3) The drug effects determined in 2 are inserted into a model of adult CMs with the same SQT1 mutation to give a prediction of drug effect in mature CMs [22]. 4) The adult CM model is converted into pseudo QT waveforms for prediction of QT segment changes in SQT1 patients under the estimated effect of the drug.

**Figure 2:**
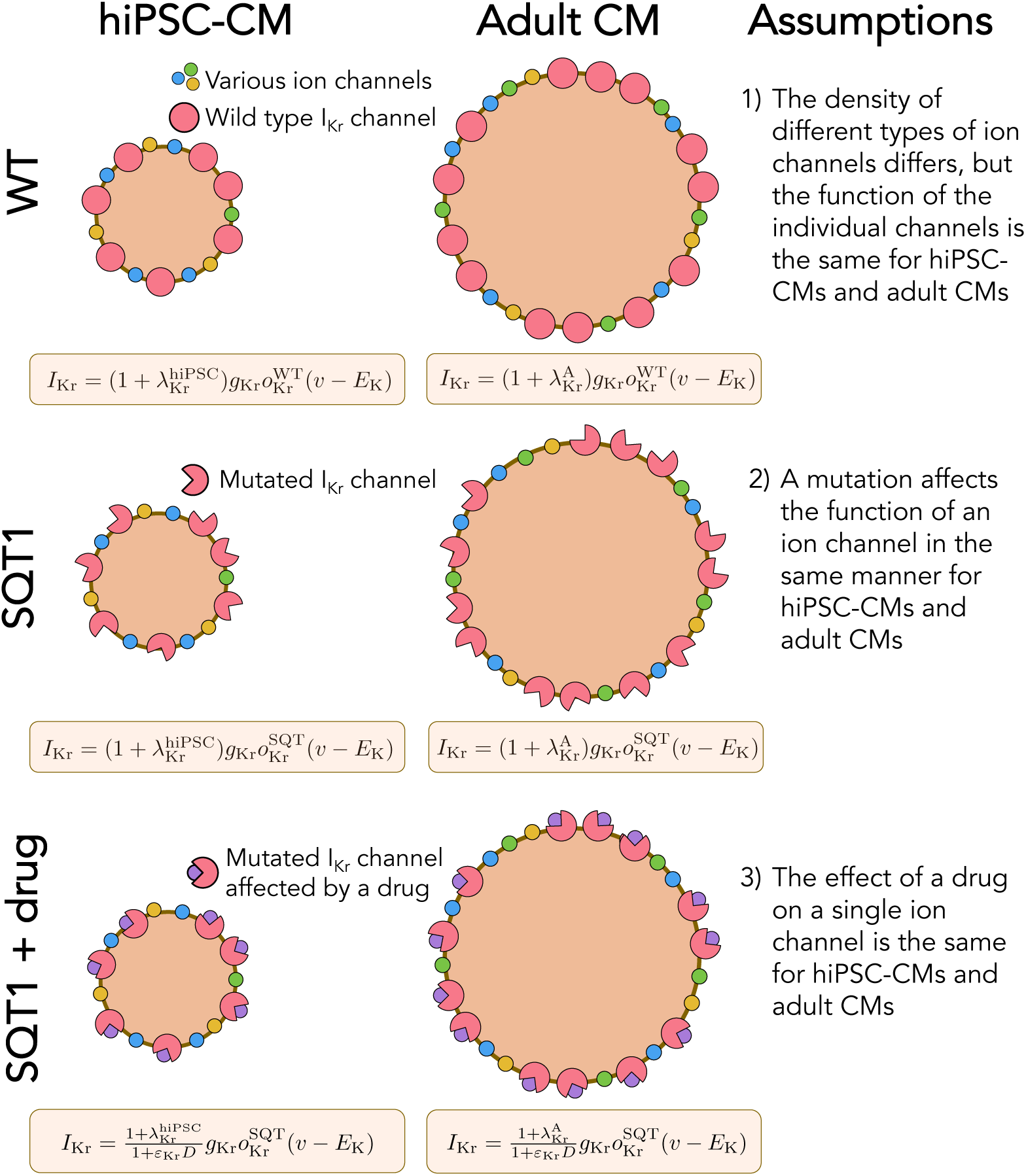
Illustration of the assumptions underlying the computational maturation approach. 1) The density of different types of ion channels (and other membrane or intracellular proteins) may differ between hiPSC-CMs and adult CMs, but the function of the individual channels is the same. In the model, the density difference is represented by the parameter *λ*. 2) The SQT1 mutation affects the individual *I*_Kr_ channels in exactly the same manner for hiPSC-CMs and adult CMs. In the model, the mutation is represented by an adjusted model for the open probability, *o*_Kr_. 3) The effect of a drug on a single protein is the same for hiPSC-CMs and adult CMs. In the model, the drug effect is represented by the parameter *ε*.

The aim of the present study is to use this computational procedure to predict drug effects for adult SQT1 CMs based on measurements of drug effects on the action potential of SQT1 hiPSC-CMs from [9, 11] and to extend these results into prediction of patient QT changes. Our overall computational pipeline is depicted in Figure 1. We consider the four drugs quinidine, ajmaline, mexilietine and ivabradine, shown to potentially be useful for SQT1 patients in [9, 11] and show predicted drug responses for the drugs on the ventricular action potential and QT interval of adult patients. We validate our pipeline using data for quinidine, where measurements of the drug effect on the QT interval have been conducted for adult patients [15]. The predicted drug response turns out to be in good agreement with the measured drug effect, indicating that the computational procedure could be useful for predicting adult drug responses from measurements of hiPSC-CMs.

## 2 Methods

In this section, we describe the methods applied in this study. We start by describing the basic modeling assumptions underlying the approaches used for computational identification of drug effects and mapping of drug effects from hiPSC-CMs to adult CMs. Then, we describe details of the applied action potential model and inversion procedure. Finally, we describe the approach used to estimate the QT interval in the adult case. Note that the majority of these methods are to a large extent based on the methods described in [23].

### 2.1 Action potential models

In this section, we describe the framework for modeling the action potentials of hiPSC-CMs and adult CMs, with and without the SQT1 mutation, and the relationship between these models. Three of the main modeling assumptions are illustrated in Figure 2, building on the approaches of [22, 23]. In the next subsections we will explain these assumptions and demonstrate how they affect the action potential modeling.

#### 2.1.1 Action potential modeling framework

In the action potential model, the membrane potential, *v* (in mV), is governed by an equation of the form

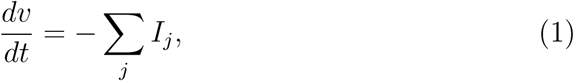

where *I*_*j*_ are different membrane current densities. These current densities can be expressed on the form

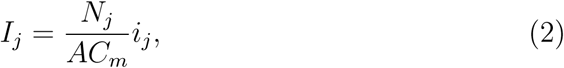

where *N*_*j*_ is the number of proteins of type *j* on the membrane, *A* is the area of the membrane, *C*_*m*_ is the specific membrane capacitance, and *i*_*j*_ is the average current through a single protein of type *j*. For currents through voltage-gated ion channels, this *i*_*j*_ is typically given on the form

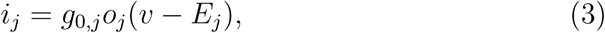

where *g*_0,*j*_ is the conductance through a single open channel, *o*_*j*_ is the open probability of the channel, and *E*_*j*_ is the equilibrium potential of the channel. In this case, it is common to introduce combined parameters of the form

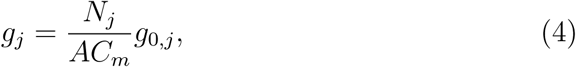

and write (2) on the form

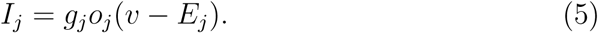

#### 2.1.2 Assumption 1: Functional invariance of wild type (WT) ion channels during maturation

A number of electrophysiological properties have been shown to differ between hiPSC-CMs and adult CMs (see e.g., [20, 21]). In addition, the electrophysiological properties of different samples of WT hiPSC-CMs often vary significantly (see e.g., [39, 40]). We assume that these differences in electrophysiological properties (both between samples of hiPSC-CMs and between adult CMs and hiPSC-CMs) are due to:

a. Differences in the geometry of the cells,
b. Differences in the number of membrane proteins, like ion channels, pumps and exchangers,
c. Differences in the number of intracellular proteins, like calcium buffers, ryanodine receptors and SERCA pumps.

However, we assume that the function of the individual membrane and intracellular proteins are the same for the different WT cases.

Considering the model (1)–(2), these assumptions imply that the density 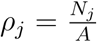 may differ between cells, but that *i*_*j*_ remains the same. Parameterizing a model to specific set of measurements can then be accomplished by adjustments of the densities represented by adjustment factors *λ*_*j*_ such that

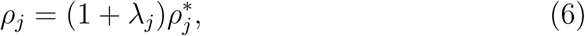

where 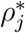 is the default density of proteins of type *j*. Incorporating these adjustment factors into the model (1), specific adult and hiPSC-CM versions of the model are given by

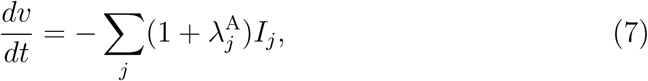

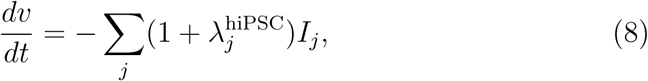

respectively, where 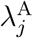 and 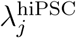 specify the protein densities in the two cases. For currents through ion channels, these adjustment factors can be incorporated by adjusting the conductances, *g*_*j*_ (see (4)–(5)). Similar adjustment factors can be set up for the density of intracellular proteins and the cell geometry, see [23]. A list of the maturation-dependent parameter values of the base model are given in Table S4 of the Supplementary Information.

#### 2.1.3 Assumption 2: Functional invariance of mutated ion channels during maturation

We assume that a mutation affects an individual ion channel in exactly the same manner for hiPSC-CMs and adult CMs. SQT1 is known to affect the *I*_Kr_ current [41, 3], and we assume that the effect on the current can be represented by adjusting the model for the open probability, *o*_Kr_ (see (3)). More specifically, we adjust the voltage dependence of the steady state inactivation gate, *x*_Kr2_ by shifting the inactivation towards more positive potentials, as described for the SQT1 mutation N588K in [3]. Because we assume that the mutation affects an individual *I*_Kr_ channel in the same manner for hiPSC-CMs and adult CMs, we apply the same adjustment of the model for *o*_Kr_ in the hiPSC and adult SQT1 cases.

#### 2.1.4 Assumption 3: Identical drug effects for identical proteins

The third assumption illustrated in Figure 2 is that since the function of a single ion channel is the same for hiPSC-CMs and adult CMs, the effect of a drug on an individual ion channel will also be identical for hiPSC-CMs and adult CMs. This means that if we are able to determine the effect of a drug on an individual channel in the hiPSC-CM case, we can find the effect of the same drug in the adult case by incorporating the same single channel drug effect in the adult version of the model.

Following [23], we use a simple IC_50_-based modeling approach to represent the effect of a drug. That is, we assume that the average single channel current through a channel *j* in the presence of the drug dose *D* is given by

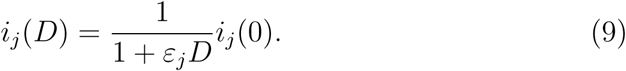

Here, *i*_*j*_(0) is the average single channel current when no drug is present and *ε*_*j*_ (in *µ*M^−1^) represents the effect of the drug, defined as

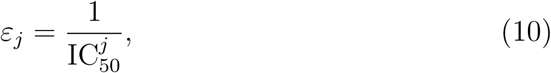

where 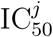 (in *µ*M) is the drug concentration that blocks the current *j* by 50%. Incorporating drug effects into the hiPSC-CM and adult CM models (7)–(8), we obtain

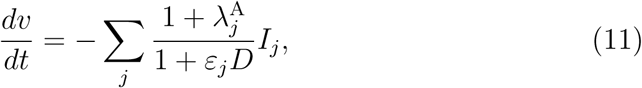

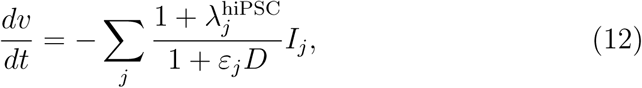

where we note that *ε*_*j*_ is the same in the two versions of the model.

In this study, we will use this modeling framework to estimate the drug effect (in the form of *ε* values) of drugs based on action potential measurements of hiPSC-CMs with the SQT1 mutation N588K. Next, we will insert the estimated *ε* values into a model for adult ventricular cells with the same mutation to estimate the effect of the drug for an adult patient.

#### 2.1.5 Base model formulation

In order to represent the action potentials of adult ventricular CMs and hiPSC-CMs (both with and without the SQT1 mutation N588K), we use a slightly modified version of the base model formulation from [23], following the form (1)–(2). Three main adjustments have been incorporated into the current version of the model. First, we have modified the steady-state values of the gating variables of the *I*_Kr_ current to better fit measured *I*_Kr_ currents in the WT and SQT1 cases from [14] (see Figure 4). Second, we have extended the temperature-dependence of the model by including Q_10_ values for a number of the currents, following [27]. This was done because we consider two different temperatures in our computations. In the adult case, we assume body temperature (310 K), while for the hiPSC-CM case, we assume room temperature (296 K) since the considered hiPSC-CM data (from [9, 11]) are obtained at room temperature. Third, we have incorporated a dynamic model for the intracellular Na^+^ concentration. This was done in order to make the frequency dependent QT interval changes more physiologically realistic (see Figure 10). The full base model formulation is found in the Supplementary Information. The system of ordinary differential equations (ODEs) defined by the base model is solved using the *ode15s* solver in Matlab.

**Figure 3:**
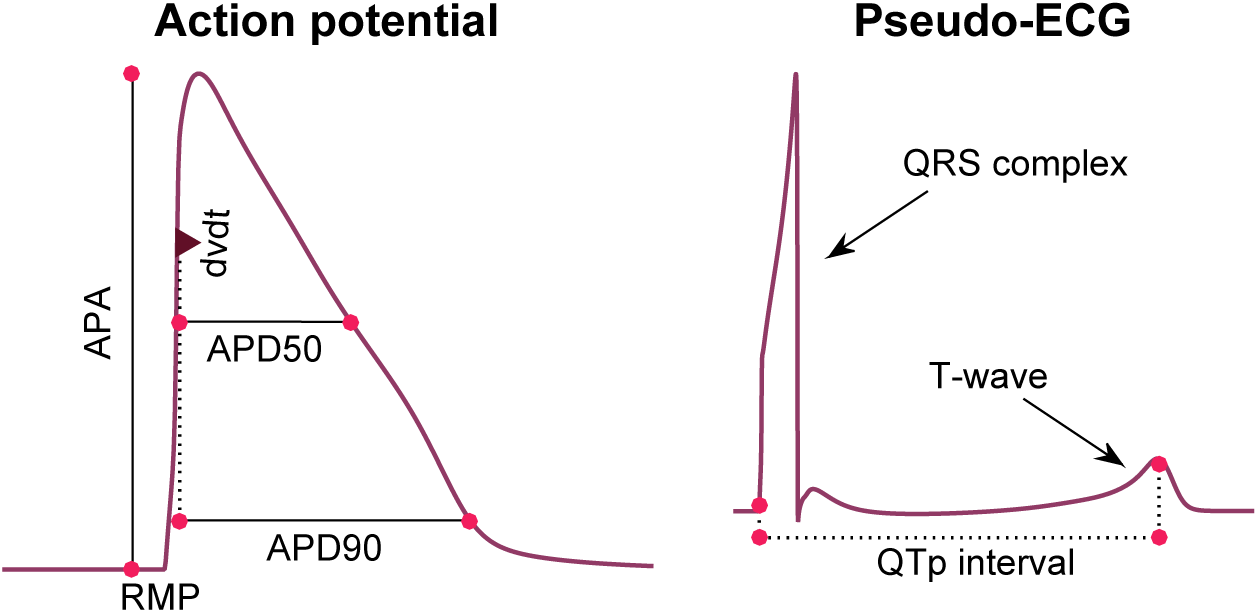
Illustration of considered biomarkers. Left: Illustration of the five biomarkers included in the cost function of the inversion procedure: The action potential durations (in ms) at 90% and 50% repolarization (APD90 and APD50, respectively), the maximal upstroke velocity (dvdt, in mV/ms), the action potential amplitude (APA, in mV) and the resting membrane potential (RMP, in mV). Right: Illustration of the QTp interval (in ms) computed from a pseudo-ECG waveform. The QTp interval is defined as the time from the onset of the QRS complex to the peak of the T-wave.

**Figure 4:**
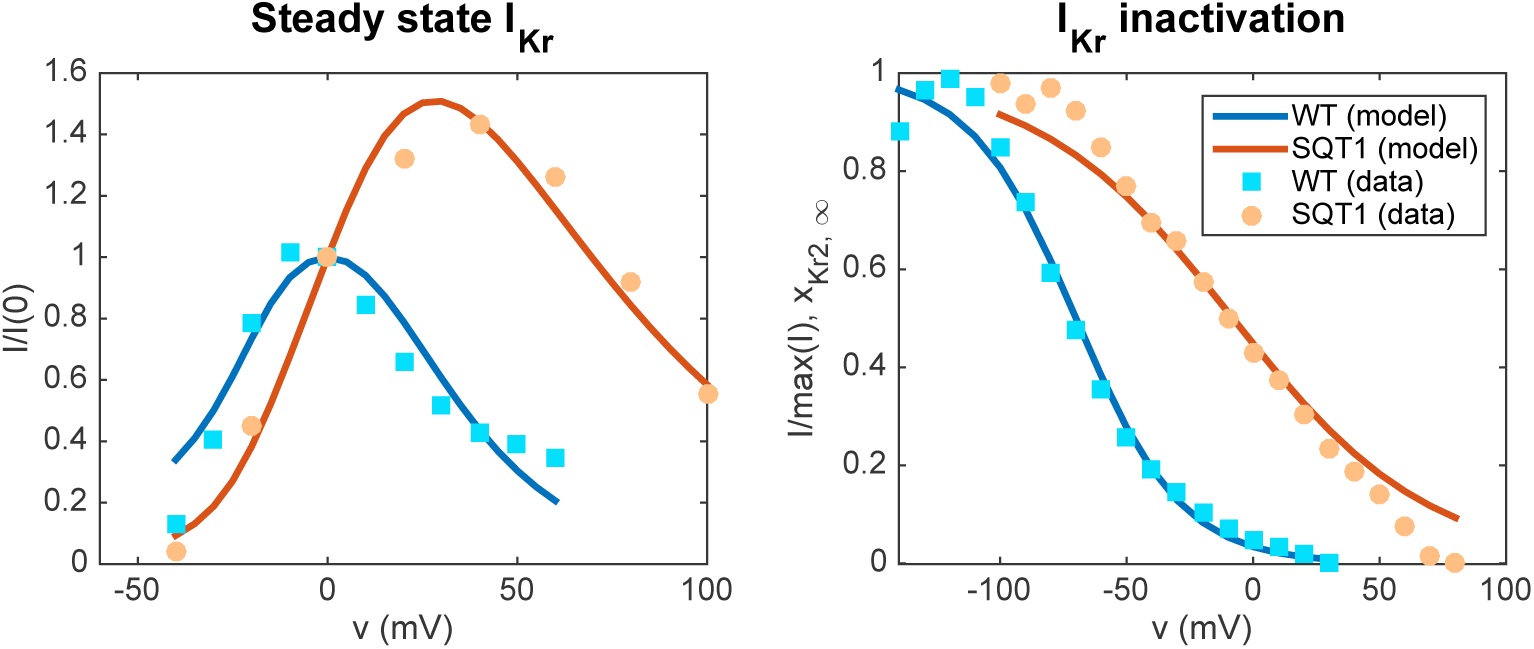
Representation of the SQT1 mutation N588K in the *I*_Kr_ model. Left panel: Steady state *I*_Kr_ currents obtained at different fixed values of the membrane potential divided by the currents obtained at *v* = 0 mV. The results obtained for the fitted WT and SQT1 *I*_Kr_ models are compared to corresponding data from [14]. Right panel: Comparison of the steady state inactivation gate in the WT and SQT1 models of *I*_Kr_ and steady state inactivation data from [14]. In the SQT1 case, the inactivation is shifted towards higher values of the membrane potential.

**Figure 5:**
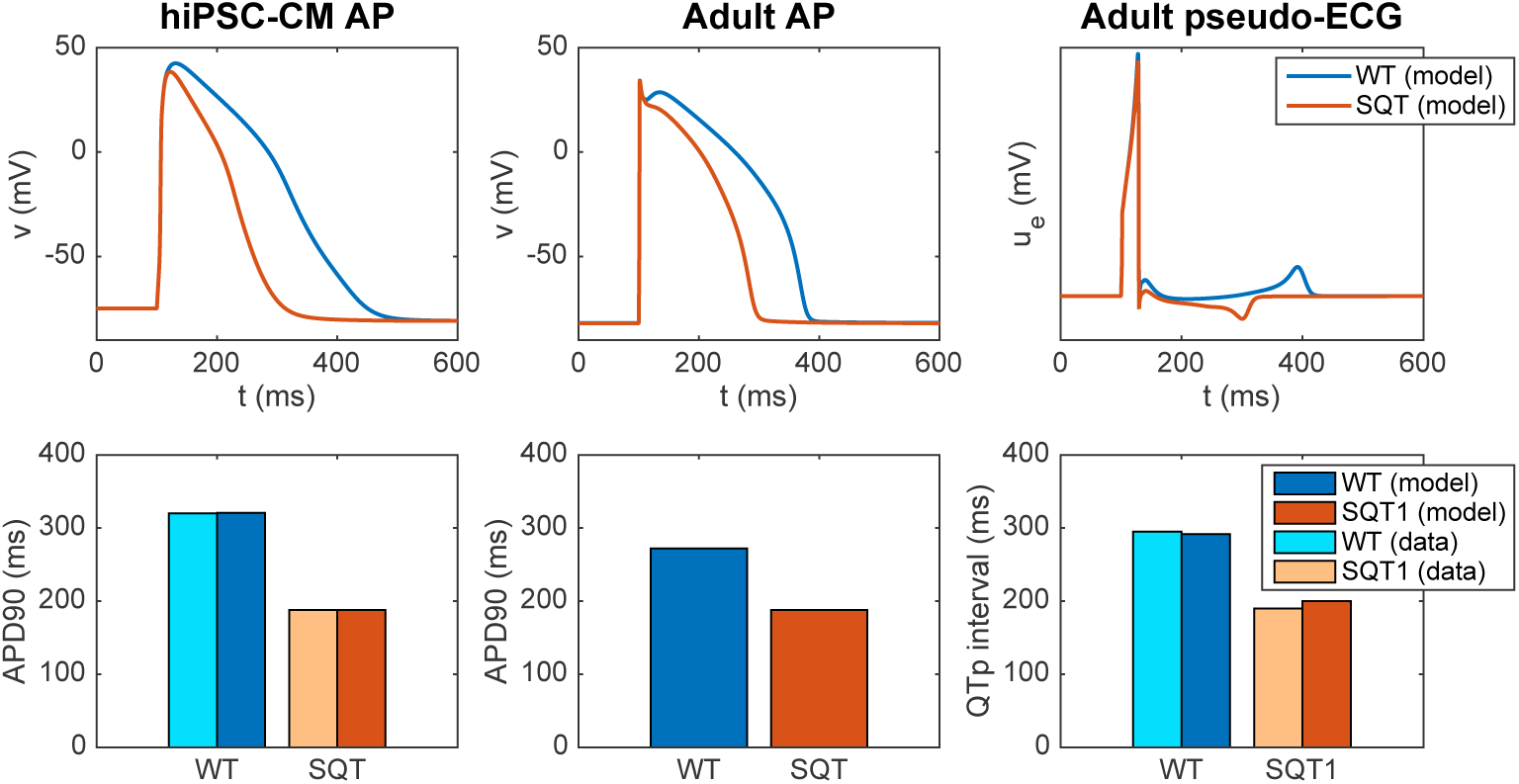
Properties of the base models for hiPSC-CMs and adult ventricular CMs in the WT and SQT1 cases. Upper panel: Comparison of the action potentials computed for the WT and SQT1 versions of the hiPSC-CM (left) and adult (center) models, in addition to the pseudo-ECG for the adult model (right). The only difference between the formulations of the WT and SQT1 models is a shift in the inactivation gate of *I*_Kr_ as illustrated in the right panel of Figure 4. Lower panel: APD90 values and QTp intervals computed using the WT and SQT1 versions of the models. The computed APD90 values for hiPSC-CMs are compared to data from [9], and the computed QTp intervals for adults are compared to data from [15].

**Figure 6:**
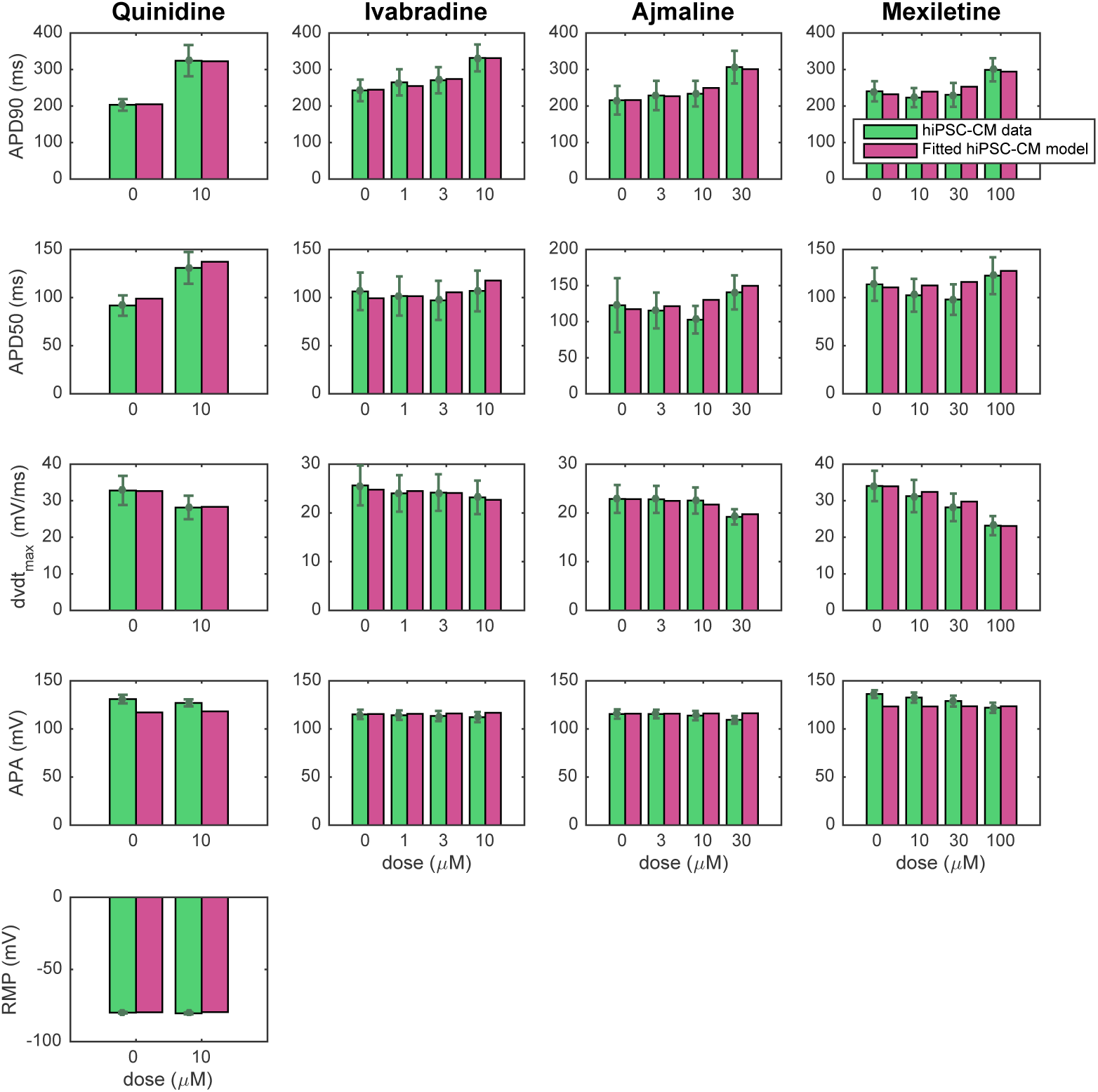
Comparison of measured and computed biomarkers. We consider the biomarkers reported in the data from [9, 11] (green) and computed from the fitted SQT1 hiPSC-CM models returned by the inversion procedure described in Section 2.2 (purple) for the drugs quinidine, ivabradine, ajmaline and mexiletine. Note that the definition of each of the biomarkers are illustrated in Figure 3 and that RMP data are only included for the quinidine case. Data are shown as the mean ± SEM (standard error of the mean).

**Figure 7:**
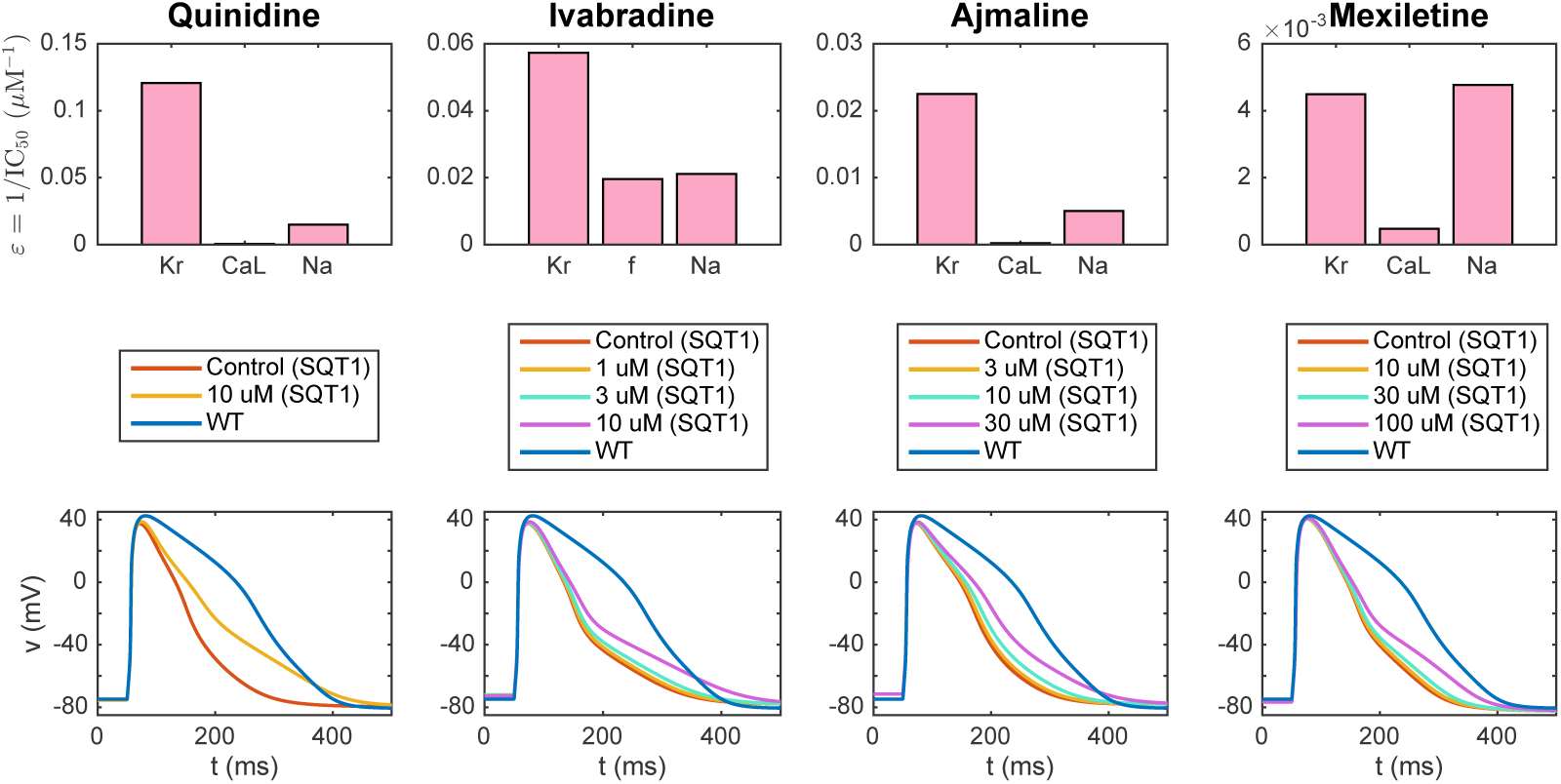
Result of the inversion procedure applied to data for the drugs quinidine, ivabradine, ajmaline and mexiletine from [9, 11]. Upper panel: Predicted drug effect on individual ion channels in the form of *ε*-values returned by the inversion procedure. Lower panel: Action potentials of the fitted SQT1 hiPSC-CM models in the control case and for doses of each drug. For comparison, we also show the action potential of the WT hiPSC-CM model.

**Figure 8:**
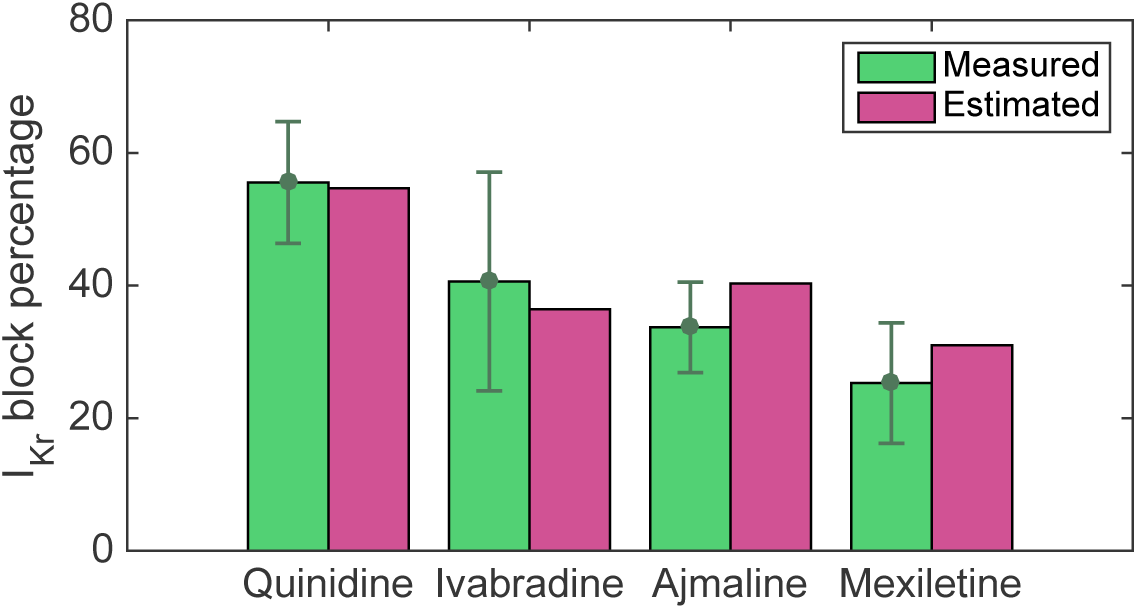
Comparison of reported and estimated block percentages for *I*_Kr_. We consider the reported block percentages for 10 *µ*M quinidine, 10 *µ*M ivabradine, 30 *µ*M ajmaline and 100 *µ*M mexiletine from [11] and the block percentages estimated based on the IC_50_ values identified in the inversion procedure. Data are shown as the mean ± SEM.

**Figure 9:**
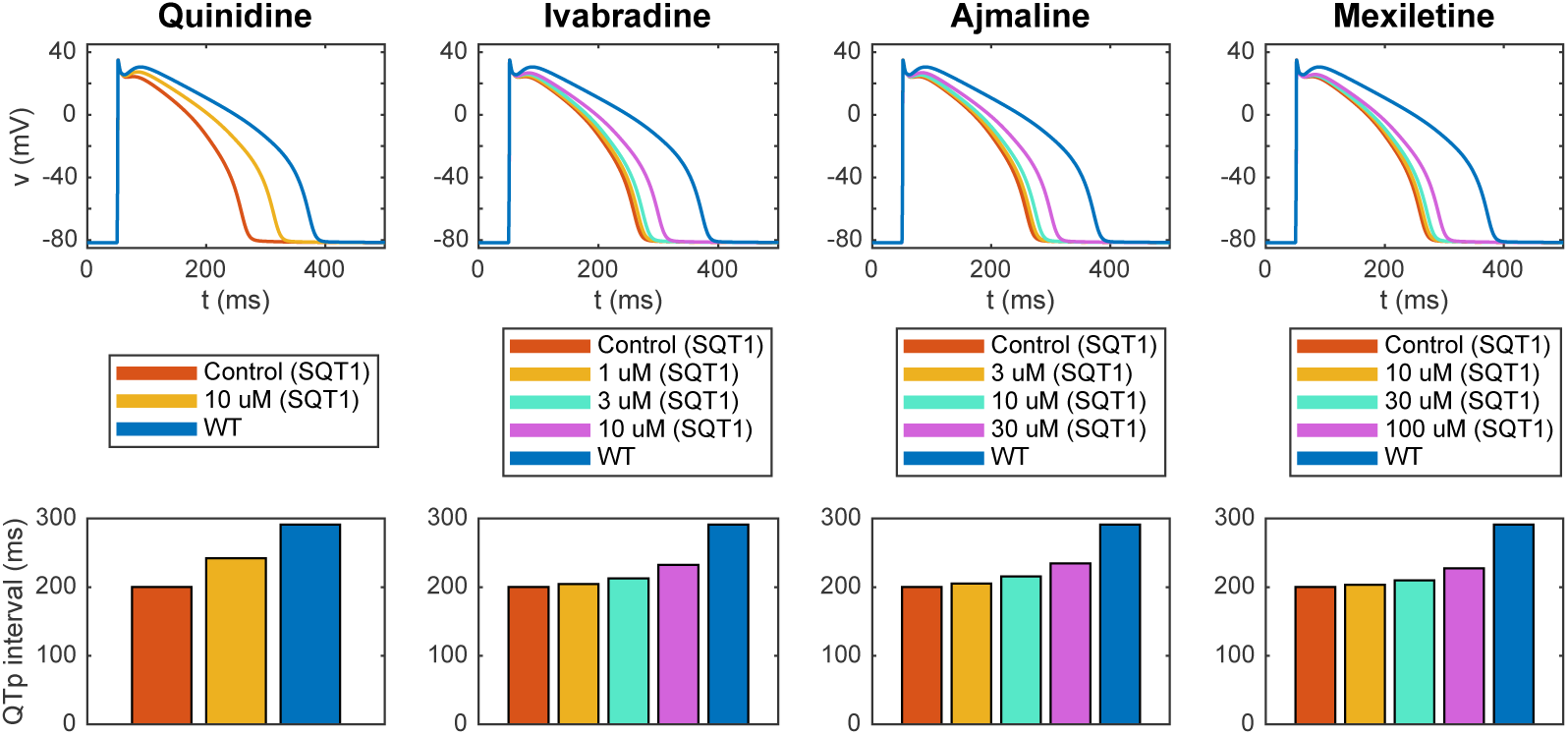
Estimated drug effects for adult patients with the SQT1 mutation. The figure displays the estimated drug effect of quinidine, ivabradine, ajmaline and mexiletine for adult patients with the SQT1 mutation (and a heart rate of 60 beats/min) based on the result of the inversion of hiPSC-CM data reported in Figure 7. Upper panel: Estimated drug effect on the adult ventricular action potential. Lower panel: Estimated drug effect on the QTp interval. See Figure 3 for an illustration of the definition of the QTp interval. Note that the blue lines represent the adult WT case.

**Figure 10:**
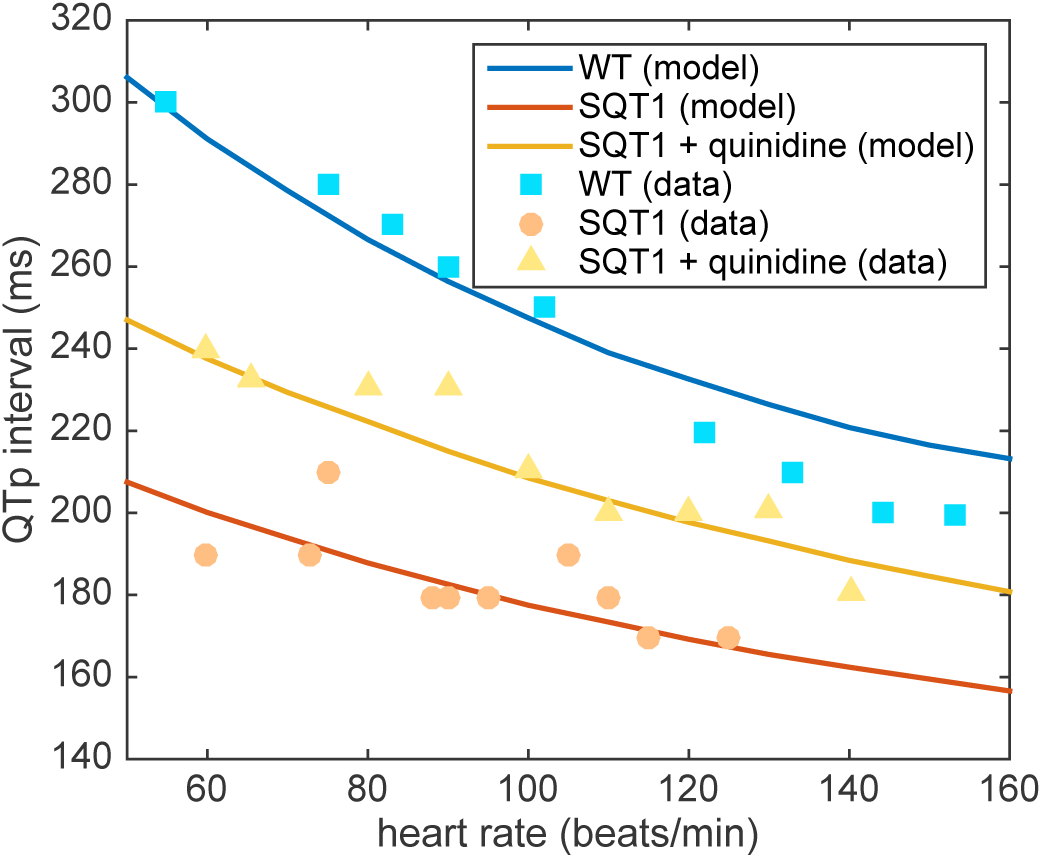
Comparison of measured and estimated adult QTp intervals. We compare the QTp intervals (see Figure 3) estimated by the computational procedure and the measured QTp intervals from [15] for different heart rates. We consider the WT case, the SQT1 case and the SQT1 case with cells exposed to 6.5 *µ*M quinidine. The drug effect estimated by the procedure is based on measured drug effects for SQT1 hiPSC-CMs from [9].

#### 2.1.6 Stimulation protocol

In the adult case, we use 1 Hz pacing unless otherwise specified. Furthermore, we run each simulation for at least 500 pacing cycles after each parameter change in order to obtain new stable solutions before recording the action potentials (and pseudo-ECGs, see Section 2.4 below).

In the hiPSC-CMs case, we use 0.2 Hz pacing, as specified in [11]. As a compromise between the need to obtain new stable solutions after each parameter change and the need for reducing the computing time of the inversion procedure, we run the simulation for 30 pacing cycles before measuring the action potential for each iteration in the continuation method used for inversion (see Section 2.2.2). However, the parameter changes between each iteration of the continuation method are expected to be quite small and we update the initial conditions between each of the 20 iterations. Therefore, the final solutions of the inversion are expected to be quite close to the steady state solutions for the applied parameters. This has also been confirmed by numerical experiments. For example, the APD90 value for the inversion of data for 10 *µ*M of quinidine was 321.4 ms using 30 pacing cycles from the states saved in the second-to-last continuation iteration. When the simulation was allowed to continue for 500 pacing cycles, the APD90 value changed by only 0.6% to 323.4 ms.

### 2.2 Inversion of action potential measurements

The inversion of data of SQT1 hiPSC-CMs exposed to various drugs is performed using the inversion procedure described in [23]. In the inversion, the adjustment factors *λ*_Kr_, *λ*_CaL_, *λ*_Na_, *λ*_K1_, *λ*_f_, *λ*_NaCa_, *λ*_NaK_, *λ*_bCl_, and *λ*_B_ for the currents *I*_Kr_, *I*_CaL_, *I*_Na_, *I*_K1_, *I*_f_, *I*_NaCa_, *I*_NaK_, *I*_bCl_, and the intracellular calcium buffers are treated as free parameters (see (8) and [23]). In addition, we assume that the drugs affect the *I*_Kr_, *I*_CaL_ or *I*_Na_ currents for quinidine, ajmaline and mexiletine and the *I*_Kr_, *I*_f_ or *I*_Na_ currents for ivabradine. Therefore, we introduce the drug parameters *ε*_Kr_, *ε*_f_ and *ε*_Na_ for ivabradine and *ε*_Kr_, *ε*_CaL_, and *ε*_Na_ for the remaining drugs.

#### 2.2.1 Cost function definition

In the inversion procedure, we wish to minimize a cost function of the form

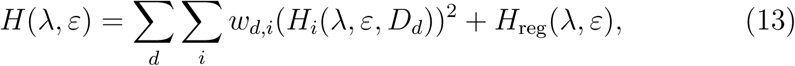

where *D*_*d*_ represent each of the drug doses included in the data set (including the control case, *D*_0_ = 0), *H*_*i*_ represent the different cost function terms included in the cost function, *w*_*d,i*_ are weights for each of the cost function terms and doses and *H*_reg_ is a regularization term (see (15) below). The cost function consists of terms of the form

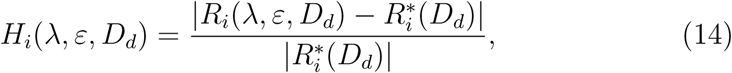

where *R*_*i*_(*λ, ε, D*_*d*_) is a biomarker computed from the solution of the model specified by *λ* and *ε* for the drug dose *D*_*d*_, and 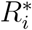(*D*_*d*_) is the corresponding measured biomarker of the data we are trying to invert.

##### Considered biomarkers

We consider each of the biomarkers, *R*_*i*_, included in the data sets from [9, 11], i.e., APD50, APD90, APA, dvdt and RMP. These biomarkers are illustrated in the left panel of Figure 3. Here, the maximal upstroke velocity, dvdt (in mV/ms), is defined as the maximum value of the derivative of the membrane potential with respect to time, the resting membrane potential, RMP (in mV), is defined as the minimum value of the membrane potential, the action potential amplitude, APA (in mV), is defined as the difference between maximum and minimum values of the membrane potential, and the APD50 and APD90 values are defined as the time (in ms) from the time point of the maximum upstroke velocity to the time point where the membrane potential first reaches a value below 50% or 90%, respectively, of the action potential amplitude.

##### Cost function weights

We use the weight 3 for *H*_APA_, the weight 6 for *H*_APD90_ and the weight 1 for the remaining terms. In addition, the weights for the control case, *w*_0,*j*_, and the weight for *H*_APD90_ for the largest dose are multiplied by the total number of doses included in the data set. The term *H*_RMP_ is only included for the drug quinidine because it was not specified in the data set for the remaining drugs [9, 11].

##### Regularization term

The regularization term, *H*_reg_(*λ, ε*) is defined as

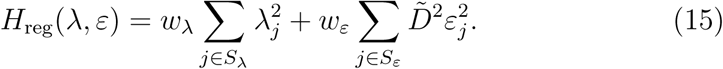

Here, the first term is defined to make the inversion procedure avoid solutions with unrealistically large perturbations of some of the currents, and the second term is defined to make the inversion select small drug perturbations if small and large perturbations result in almost indistinguishable solutions (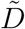 is the median of the considered drug doses of the data set). In our computations, we set the weights for the terms to *w*_*λ*_ = 1 and *w*_*ε*_ = 0.001. We let the set *S*_*ε*_ consist of all the currents we assume may be affected by the drug and the set *S*_*λ*_ consist of the *I*_Kr_ and *I*_CaL_ currents.

#### 2.2.2 Minimization method

We apply the continuation-based minimization procedure from [23] to minimize the cost function (13). We use 20 continuation iterations with 100 or 200 randomly chosen initial guesses for the first fifteen and the last five iterations, respectively. The initial guesses for *λ* are chosen within 20% above or below the optimal values from the previous iteration, and the initial guesses for *ε*_*m*_ are chosen within [*ε*_*m*−1_*/*5, 5*ε*_*m*−1_], where *ε*_*m*−1_ is the optimal *ε* from the previous iteration. From these initial guesses we run 30 or 60 iterations of the Nelder-Mead algorithm [42] for the first fifteen and the last five iterations, respectively.

### 2.3 Inversion of *I*_Kr_ measurements

In order to represent the WT and SQT1 *I*_Kr_ currents, parameters of the model for the *I*_Kr_ open probability are fitted to data of WT and SQT1 *I*_Kr_ currents from [14]. In this data, it is revealed that the steady state inactivation of the current is shifted towards higher potentials in the SQT1 (N588K) case compared to the WT case (see the right panel of Figure 4). Therefore, we assume that the mutation can be represented in the model by an adjusted model for the steady state value of the inactivation gate, *x*_Kr2_. To find a model matching the data as well as possible, we introduce the six free parameters *c*_1_ − *c*_6_ in the steady state activation and inactivation gates of the *I*_Kr_ current on the form:

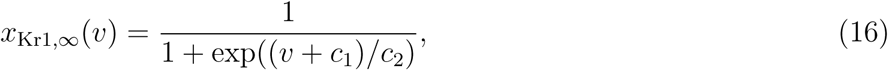

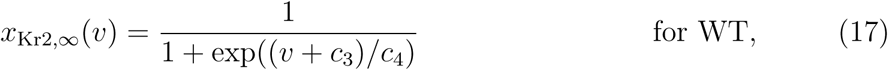

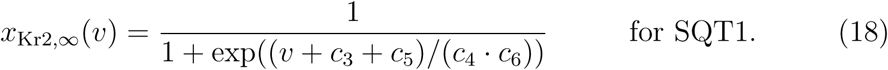

See the Supplementary Information for a description of the full *I*_Kr_ model formulation. To find optimal values of the six parameters *c*_1_ − *c*_6_, we run simulations of the voltage clamp protocol applied in [14]. That is, we first fix the membrane potential at -80 mV and run a simulation to steady state, then we increase the membrane potential to a value between -50 mV and 100 mV and compare the current after 2 seconds of simulation to the corresponding current reported in [14] (see left panel of Figure 4). In addition, we compare the steady state values of the inactivation gate *x*_Kr2_ directly to the steady state inactivation measurements reported in [14].

In the simulations trying to adjust the *I*_Kr_ model to measurements from [14], we adjust the temperature and intracellular and extracellular potassium concentrations in the model to match those reported for the experiments, that is *T* = 37 °C, [K^+^]_*i*_ = 130 mM, [K^+^]_*e*_ = 4 mM.

We find the optimal parameters *c* = (*c*_1_, …, *c*_6_) by minimizing a cost function of the form

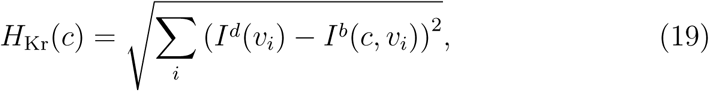

using the Nelder-Mead algorithm [42]. Here, *v*_*i*_ are each of the values of the membrane potential considered in the data set, *I*^*d*^(*v*_*i*_) are each of the measured data points for relative *I*_Kr_ currents, and *I*^*b*^(*c, v*_*i*_) are the corresponding currents computed for the model specified by the parameters *c*.

### 2.4 Computation of pseudo-ECG and QT interval

In order to estimate drug effects on adult QT intervals, we apply a simple pseudo-ECG calculation applied in a number of earlier studies (e.g., [43, 44, 45, 46, 47]). The calculation follows a two-step procedure.

**Step 1** The membrane potential and membrane currents along a strand of cylindrical cells are computed using the cable equation as explained i detail in [35]. In this step, cells are connected to each other by gap junctions, each cell is assumed to be isopotential, and the extracellular potential is assumed to be constant in space. The membrane potential *v*_*k*_ in each cell *k* is modeled by

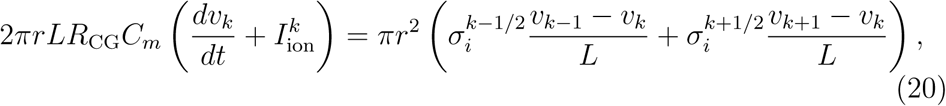

where *r* is the cell radius, *L* is the cell length, *R*_CG_ is the ratio between the capacitive and the geometrical cell areas (see e.g., [35]), *C*_*m*_ is the specific cell capacitance and 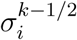 is the averaged intracellular conductivity between cells *k* − 1 and *k*. This averaged intracellular conductivity is given by

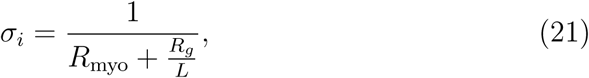

where *R*_myo_ is the myoplasmic resistance and *R*_*g*_ is the gap junction resistance. Furthermore, 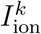 is the sum of the ionic current densities of the cell, modeled by the base model (i.e.,Σ_*j*_ *I*_*j*_ in (1)).

The ODE system defined by (20) is solved using the *ode15s* solver in Matlab, and the solution is used to compute the membrane currents

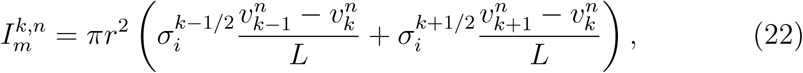

originating from each cell *i* at each time point *n*.

**Step 2** The extracellular potential originating from the cell strand is computed for each time step *n* using the so-called point-source approximation (see e.g., [43, 48])

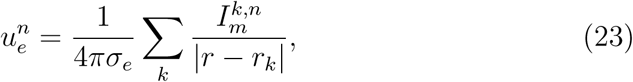

where *σ*_*e*_ is the extracellular conductivity and |*r* − *r*_*k*_| is the Euclidean distance between the point at which we are measuring the extracellular potential, *r*, and the center of cell number *k, r*_*k*_.

#### Parameter values

The parameters of the pseudo-ECG simulations are based on the parameters of earlier computations of pseudo-ECGs (e.g., [43, 46, 35]). We consider a cell stand of 100 cells of length *L* = 150 *µ*m and radius *r* = 10 *µ*m. We let the cell strand exhibit transmural heterogeneity of ion channel density, with an endocardial region consisting of the first 25 cells, a midmyocardial region for the next 35 cells and an epicardial region in the last 40 cells. The default adult base model defines the ion channel density in the epicardial region. In the endocardial region, the *I*_to_ and *I*_Ks_ channel densities are reduced to 1% and 31%, respectively, compared to the epicardial region. Similarly, in the midmyocardial region, the *I*_to_ and *I*_Ks_ channel densities are reduced to 85% and 11%, respectively, (compared to the epicardial region).

Furthermore, we set *C*_*m*_ = 1 *µ*F/cm^2^, *R*_CG_ = 2, *R*_myo_ = 0.15 kΩcm, and *R*_*g*_ = 0.0015 kΩcm^2^, resulting in a conduction velocity along the cell strand of approximately 52 cm/s, close to experimentally measured conduction velocities from literature (∼ 50 cm/s [49]). Stimulation is applied for the first two cells of the endocardial region, and the extracellular potential is measured 2 cm from the end of the epicardial region. Because we in this study only are interested in the time course of the ECG and not the amplitude in mV, we do not assign a specific value for *σ*_*e*_ and report the computed pseudo-ECG without numbers on the *u*_*e*_-axis.

#### Definition of the QT interval

After computing a pseudo-ECG using the described approach, the QT interval is computed from the ECG waveforms. The QT interval is often defined as the time from the start of the QRS interval to the end of the T-wave (see Figure 3). However, in the study [15], which we will use for comparison to the computational results, an alternative definition of the so-called QTp interval is applied, defined as the time from the start of the QRS complex to the peak of the T-wave, as illustrated in the right panel of Figure 3. We will therefore use this definition in this study. Note that for the SQT1 case, the computed T-wave is inverted to have the opposite sign of the QRS complex (see Figure 5). In that case, we define the peak of the T-wave as the time point when the minimum, i.e. the maximum absolute deviation of the T-wave, is reached.

## 3 Results

In this section, we describe the results of our applications of the above mentioned methods. First, we set up the default hiPSC-CM and adult base models for WT and SQT1 based on measurements of WT and SQT1 *I*_Kr_ currents from [14]. Next, we apply the inversion procedure to identify the effect of four drugs on individual ion channels based on action potential measurements of SQT1 hiPSC-CMs from [9, 11]. Finally, we estimate the corresponding drug effect for adult patients with short QT syndrome using these identified drug effects.

### 3.1 Computational representation of the SQT1 mutation

In order to represent the SQT1 mutation N588K in the model, we first adjust the model of the open probability of the *I*_Kr_ current to data for WT and SQT1 *I*_Kr_ currents from [14] as described in Section 2.3. The optimal values for the model given on the form (16)–(18) returned by the inversion procedure are *c*_1_ = −2.7, *c*_2_ = −15.3, *c*_3_ = 70, *c*_4_ = 20.9, *c*_5_ = −62, *c*_6_ = 1.85. In Figure 4, the fitted model solutions of *I*_Kr_ in the WT and SQT1 cases are compared to the data from [14]. In the right panel, we observe that the inactivation is shifted towards higher values of the membrane potential in the SQT1 case.

The WT and SQT1 versions of the *I*_Kr_ model fitted to measurements from [14] are inserted into the base model formulation. The parameters, *λ*^hiPSC^, for the hiPSC-CM versions of the model are then fitted to action potential measurements of WT and SQT1 hiPSC-CMs from [9]. In addition, the parameters of adult versions of the base model are fitted to information about the QTp interval for adults with and without the SQT1 mutation from [15]. These adjustments are made by hand-tuning the parameters of the default hiPSC-CM and adult versions of the base model from [23].

The action potentials of the resulting models are plotted in the upper panel of Figure 5, along with the computed pseudo-ECG for the adult case. The pseudo-ECG is computed using the approach described in Section 2.4. Note that the only difference between the WT and SQT1 versions of the models is a difference in the steady state inactivation gate of the *I*_Kr_ current (see (18)), and that the remaining parameters of the models are the same in the WT and SQT1 cases. Note also that the adjustment of the *I*_Kr_ steady state inactivation gate is exactly the same in the hiPSC-CM and adult versions of the model (following assumption 2 in Figure 2), but that the geometry of the cell and the density of different types of membrane and intracellular proteins are different between the models for hiPSC-CMs and adult CMs (following assumption 1 in Figure 2).

In the lower panel of Figure 5, we report the APD90 values of the WT and SQT1 models in the hiPSC-CM and adult cases. In addition, we report the QTp interval computed from the pseudo-ECG (see Figure 3). In the hiPSC-CM case, we compare the APD90 values of the model to the corresponding values reported in [9]. The APD90 value is reduced from 321 ms in the WT case to 188 ms in the SQT1 case, in good agreement with the values reported in [9]. In the adult case, the APD90 value is similarly reduced from 272 ms to 188 ms. Furthermore, the QTp interval is reduced from 292 ms in the WT case to 200 ms in the SQT1 case, close to reported QTp values from [15].

### 3.2 Computational identification of drug response from membrane potential measurements of SQT1 hiPSC-CMs

In [9, 11], data are reported from measurements of action potential biomarkers for hiPSC-CMs with the SQT1 mutation N588K exposed to drugs attempting to make the properties of the SQT1 cells more similar to WT cells. In this paper, we wish to use this data to estimate the corresponding drug responses for adult SQT1 patients. In order to make these predictions, we first need to identify the effect of the drugs on individual ion channels in the hiPSC-CM case, based on the data provided in [9, 11]. These drug effects are predicted using the inversion procedure described in Section 2.2 using the SQT1 hiPSC-CM model from Figure 5 as a starting point for the inversion.

Figure 6 shows how well the biomarkers of the fitted SQT1 hiPSC-CM model match the corresponding biomarkers reported in the data. We observe that the biomarkers of the fitted models are quite similar to the values of the data. In particular, the model seem to capture the APD90 increase resulting from the drugs quite well. In the lower panel of Figure 7, the action potentials of the SQT1 hiPSC-CM models fitted to the data are plotted for the control case and for each of the drug doses, along with the action potential of the default base model for WT hiPSC-CM. We observe that each of the drugs increase the action potential duration, but not enough to fully recapture the WT hiPSC-CM action potential length.

The upper panel of Figure 7 reports the drug effects identified by the inversion procedure in the form of *ε* values. Recall that *ε*_*i*_ is defined as 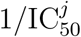, where 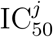 is the drug concentration that blocks the current *j* by 50%. In other words, a high *ε* value corresponds to a significant drug effect. We observe that the inversion procedure predicts that all four drugs have a significant effect on the *I*_Kr_ current.

In [11], measured drug effects on SQT1 *I*_Kr_ currents are reported for the maximum dose of each of the considered drugs. In order to get an impression of the accuracy of the IC_50_ values estimated by the inversion procedure, we compare the block percentages reported in [11] to the corresponding block percentages estimated by the computational procedure. The results are displayed in Figure 8. We observe that the estimated block percentage is very similar to the measured valued for the drugs quinidine and ivabradine. For ajmaline and mexiletine, the computational procedure seems to predict a slightly larger block percentage than what was found in the measurements. However, all identified block percentages are within the standard error of the mean reported for the measurements.

### 3.3 Estimation of drug response for adult SQT1 patients

Following assumption 3 in Figure 2, we assume that the effect of a drug on a single ion channel is the same for hiPSC-CMs and adult CMs. Using this assumption, we can estimate the effect of the drugs quinidine, ivabradine, ajmaline and mexiletine for adult SQT1 CMs by inserting the *ε* values identified in Figure 7 into the adult SQT1 base model. In Figure 9, we report the resulting estimated adult drug effects. In the upper panel, we show the ventricular action potentials for the SQT1 case with no drug and with each of the considered drug doses present. In addition, we show the action potential in the WT case. The lower panel of Figure 9 displays the adult QTp intervals computed from pseudo-ECG simulations as explained in Section 2.4. We observe that the drugs increase the QTp interval for the SQT1 case to be more similar to the WT QTp interval. For 10 *µ*M of quinidine, the QTp interval is estimated to increase from 200 ms to 242 ms. Similarly, the QTp interval is predicted to increase to 232 ms for 10 *µ*M of ivabradine, 235 ms for 30 *µ*M of ajmaline and 227 ms for 100 *µ*M of mexiletine. For comparison, the WT QTp interval is 292 ms.

The effect of the drug quinidine on the QTp interval of adult SQT1 patients is reported for a number of different heart rates in [15]. In Figure 10, we compare the predicted adult drug response of quinidine from our computational procedure to the measured responses in [15]. In the study [15], mean serum concentration was reported as 2.1 mg/L. Applying the molecular weight of quinidine of 324.4 g/mol [50], we consider the drug concentration 6.5 *µ*M in the computational model. Moreover, we use the *ε* values identified based on measurements of SQT1 hiPSC-CMs from [9] (see Figure 7). The QTp intervals are computed for different pacing frequencies using the pseudo-ECG approach described in Section 2.4. In Figure 10, we observe that the QTp intervals predicted by the model for different heart rates fit well with the measured data in both the WT case, in the SQT1 case and in the SQT1 case with quinidine.

## 4 Discussion

In this paper, we have applied the computational procedure introduced in [22, 23] to data of hiPSC-CMs derived from an SQT1 patient, found in [9, 11]. In this procedure, we first identified the drug response on individual ion channels based on measurements of drug effects for SQT1 hiPSC-CMs and then estimated the drug effect for an adult SQT1 patient by inserting these single channel drug effects into a model for adult cells.

### 4.1 Computational representation of the SQT mutation

Short QT syndrome is characterized by an abnormally short duration of the QT interval, and several different genetic mutations have proven to give rise to different subtypes of the syndrome [1, 2, 3]. SQT1–SQT3 are caused by gain of function mutations in the potassium channels responsible for the *I*_Kr_, *I*_Ks_ and *I*_K1_ currents, respectively, while SQT4–SQT6 are associated with loss of function mutations in genes encoding calcium channels. Furthermore, SQT7 and SQT8 have been characterized by a loss of function mutation in a gene encoding sodium channels and a mutation in a gene encoding the cardiac Cl/HCO_3_ exchangers AE3, respectively [51]. Because of the different effects of different mutations, it is likely that pharmacological treatment of SQT syndrome must be tailored to the individual subtypes. Furthermore, in order to study the effects of mutations computationally, different mathematical representations are needed for the mechanistic causes of the shortening.

In this study, we have considered the SQT1 subtype, caused by a mutation (N588K) leading to increased *I*_Kr_ currents. Based on measurements of WT and mutated *I*_Kr_ currents [14], we represented the mutation by shifting the inactivation of the channels towards more positive potentials (see (18) and Figure 4). This computational representation of the mutation follows the same lines as [52, 53, 46], which used similar adjustments of the model for the *I*_Kr_, *I*_Ks_ and *I*_K1_ open probabilities to represent the SQT1, SQT2 and SQT3 subtypes, respectively.

In the base model used in this study (see the Supplementary Information), as well as in other commonly used action potential models (e.g., [26, 27, 54, 30]), the open probability of voltage-gated ion channels are governed by gating variables following the Hodgkin-Huxley formalism. However, more detailed and versatile models for the open probability have been introduced in the form of more complicated Markov models (see e.g., [55, 56] for a comparison of the two formalisms). Such Markov models have been suggested to give a more realistic representation of both the effect of mutations and the effect of drugs (see e.g., [57, 58, 59]), and was applied in computational studies of SQT1 in [45, 60, 61] and SQT3 in [62]. On the other hand, complex Markov models are associated with a significant increase in the number of free parameters to determine in the models, and we have therefore chosen to use a more simplified Hodgkin-Huxley formalism in this study.

The WT and SQT1 versions of the *I*_Kr_ current were inserted into the base models for hiPSC-CMs and adult CMs to define WT and SQT1 versions of these action potential models. The resulting computed action potentials displayed a clear action potential shortening for the SQT1 case both for hiPSC-CMs and adult CMs (see Figure 5). The computed pseudo-ECGs for the adult case also displayed a clear reduction in the duration of the QT interval for SQT1. Furthermore, the APD90 values of the hiPSC-CM model and the QT intervals of the adult model appeared to fit well with measured data for WT and SQT1 from [9, 15].

### 4.2 Computational identification of drug effects from membrane potential measurements of SQT1 hiPSC-CMs

Because of the possibly lethal risks associated with SQT syndrome, there is an urgent need for efficient therapies, including pharmacological treatment. In [9, 11], a number of drug candidates were tested on hiPSC-CMs generated from an SQT1 patient, and the drugs quinidine, ivabradine, ajmailine and mexiletine seemed to be the most promising for increasing the action potential duration of the SQT1 hiPSC-CMs. Other drugs shown to block WT *I*_Kr_ (sotalol, ranolazine, flecainide, amiodarone) did not seem to work as well for hiPSC-CMs affected by the SQT1 mutation. This highlights the need for mutation-specific investigations of drug effects. Here we have tried to extend such investigations computationally, and have used the measurements from [9, 11] to estimate the effect of quinidine, ivabradine, ajmailine and mexiletine for adult SQT1 patients using the computational procedure introduced in [22, 23].

#### 4.2.1 Identifiability of model parameters

In [22, 23], drug effects were identified in an inversion procedure based on optical measurements of the membrane potential and cytosolic calcium concentration of hiPSC-CMs in a microphysiological system [63]. In the present study, we have used a different type of data in the inversion procedure, i.e., membrane potential measurements from patch-clamp recordings. This alternative data type could have both advantages and disadvantages compared to the optical measurements used in [22, 23]. A disadvantage is that we only have measurements of the membrane potential and not any information about the calcium transient. Information about the calcium transient has been shown to potentially improve the identification of parameters in mathematical action potential models [22, 64, 65]. However, an advantage of the patch-clamp recordings is that we can obtain information about the actual value of the membrane potential instead of just relative pixel intensities from optical measurements of voltage sensitive dyes. Therefore, we can get information about the resting membrane potential, the action potential amplitude and the upstroke velocity, which could all be useful for the identification of parameters. For example, the upstroke velocity is difficult to obtain from optical measurements of the action potential, which makes it is very hard to identify drug effects on the fast sodium current, *I*_Na_, unless additional data, e.g. extracellular measurements are included (see e.g., [66]).

With information about the upstroke velocity, it should be possible to identify *I*_Na_. However, full identification of all model parameters based on membrane potential measurements is very challenging, and in some cases several combinations of parameter values result in virtually identical action potentials (see e.g., [67, 65]). In order to investigate which currents were identifiable in the SQT1 hiPSC-CM base model, we applied the singular value decomposition approach from [68]. The result of this analysis is given in the Supplementary Information. In short, the analysis reveals that the *I*_Na_, *I*_CaL_ and *I*_Kr_ currents are the most identifiable currents of the model. These are the three main currents investigated for drug effects in our study. In addition, we estimate drug effects on *I*_f_ for the *I*_f_ -blocking drug ivabradine. However, we note that the identifiability of this current is not very high and that the estimated drug effect for *I*_f_ consequently is associated with a large degree of uncertainty.

#### 4.2.2 Identified drug effects

Using the computational inversion procedure described in Section 2.2, we estimated drug effects on major ion channels based on action potential measurements of SQT1 hiPSC-CMs. For quinidine, ajmaline and mexiletine, we considered drug effects on the currents *I*_Kr_, *I*_CaL_ and *I*_Na_, and for ivabradine, we considered drug effects on the currents *I*_Kr_, *I*_f_ and *I*_Na_, because these currents have been shown to be affected by ivabradine [69, 70, 71].

Each of the considered drugs had a considerable effect of the *I*_Kr_ current, with predicted IC_50_ values of 8.3 *µ*M for quinidine, 17 *µ*M for ivabradine, 44 *µ*M for ajmaline and 223 *µ*M for mexiletine (see Figure 7). The effect of a drug can be very different for WT *I*_Kr_ channels compared to that of mutated *I*_Kr_ channels (see e.g., [72, 73]). For instance, in [72], the IC_50_ value of quinidine was found to be 0.6 *µ*M for WT *I*_Kr_ channels and 2.2 *µ*M for the SQT1 mutation N588K. Consequently, it is not reasonable to compare the IC_50_ values identified for the SQT1 case to IC_50_ values found in WT *I*_Kr_ experiments. However, we observe that the identified IC_50_ value for quinidine is quite close to the IC_50_ value reported for the SQT1 case in [72] (8.3 *µ*M vs 2.2 *µ*M). Furthermore, the predicted block percentage for 10 *µ*M of quinidine is very close to the measured block percentage reported in [11] (see Figure 8). The block percentages for the remaining drugs are also quite close to the percentages reported in [11], indicating reasonable identified IC_50_ values for SQT1 *I*_Kr_. For ivabradine, the identified IC_50_ value for *I*_Kr_ (17 *µ*M) is also in good agreement with the IC_50_ value of 10.3 *µ*M reported for *I*_Kr_ affected by the SQT1 (N588K) mutation in [69].

In addition, the inversion procedure identified a significant block of *I*_Na_ for the drug mexiletine (see Figure 7). This is reasonable, since mexiletine is known as an *I*_Na_ blocker [74]. However, the identified IC_50_ value of 210 *µ*M is larger than IC_50_ values found in literature (7.6 *µ*M – 43 *µ*M) [75, 76]. Moreover, ivabradine is an *I*_f_ blocker with an IC_50_ value found to be about 2 *µ*M in [70]. The computational procedure is not able to correctly identify this effect, and underestimates the drug effect to an IC_50_ value of about 50 *µ*M. This discrepancy is not unexpected, since the identifiability of *I*_f_ is predicted to be low (see Figure S1 in the Supplementary Information). The IC_50_ value for *I*_Na_ block of ivabradine, on the other hand, was estimated to 47 *µ*M, in good agreement with the measured IC_50_ value of 30 *µ*M reported in [71]. Possible effects of ajmaline and quinidine on *I*_CaL_ and *I*_Na_ and of mexiletine on *I*_CaL_ are underestimated by the inversion procedure compared to the IC_50_ values reported in [76]. This could indicate that the biomarkers considered in the inversion procedure (see Figures 3 and 6) are not enough to properly characterize these less pronounced drug effects or that the applied base model formulation or simplified drug modeling are not sufficient to represent these effects. It also could be that the regularization term used for the *ε* values can cause underestimation due to its pressure on finding the minimum effect in the chosen fit. It should also be noted that IC_50_ values identified in different experiments tend to vary considerably.

### 4.3 Computational maturation of drug response

Using the single channel drug effects identified from membrane potential measurements of hiPSC-CM, we estimated the drug effects in the adult case based on the three assumptions explained in Figure 2. First, we assumed that the density of different types of ion channels (and other membrane and intracellular proteins) differ between hiPSC-CMs and adult CMs, but that the function of a single channel is the same for adult CMs and hiPSC-CMs. This means that a set of parameters representing the density of different types of proteins, e.g. ion channel conductances (see (4)), are different between the hiPSC-CM and adult CM models, but that the models for, e.g., the open probability of the channels are the same in both versions of the model. Second, we assumed that a mutation affects the function of an ion channel in exactly the same manner for hiPSC-CMs and adult CMs. Therefore, we used the same adjustment of the open probability of the *I*_Kr_ current (see Figure 4) to represent the SQT1 mutation N588K for both hiPSC-CMs and adult CMs (see Figure 5). Third, we assumed that a drug affects a single ion channel in exactly the same manner for hiPSC-CMs and adult CMs, and we could therefore estimate the drug effect in the adult case based on the drug effect identified for hiPSC-CMs.

This modeling framework was used in [22, 23] to predict side effects of drugs for adult WT CMs based on measurements of WT hiPSC-CMs. However, the procedure can also take the effect of mutations into account (as specified in the second assumption of Figure 2), and in the present study, we have used the procedure to estimate drug effects on the ventricular action potential and the QT interval of adult SQT1 patients based on measurements of hiPSC-CMs generated from an SQT1 patient.

#### 4.3.1 Estimation of the QT interval

Because the effect of one of the drugs considered in this study has been investigated for adult SQT1 patients in the form of measured QT intervals of patients’ ECGs, we wished to be able to compare the drug responses predicted by our computational procedure to these measured QT intervals. However, the mathematical base model from [23] only represents the adult ventricular action potential and cannot be directly used to compute the adult QT interval. On the other hand, it is generally recognized that there is a clear link between the duration of the ventricular action potential and the QT interval of a patient’s ECG. For example, in [77], a simple linear relationship between the APD90 value and the QT interval was observed. Conversely, in the computational study [78], it was shown that there was not a direct linear realtionship between drug effects on the action potential duration and drug effects on the QT interval. For example, drugs blocking the *I*_Na_ current gave rise to prolongations of the computed QT intervals, even though no prolongation was observed for the action potential duration. In this study, we have chosen to predict the QT interval by computing a pseudo-ECG using a very simple mathematical representation, as applied in several previous studies of drug effects on the QT interval (e.g., [43, 46, 47]). The applied method appeared to give rise to reasonable pseudo-ECG waveforms and QT intervals (see e.g., Figure 5), but it is a very simplified representation of the dynamics underlying the ECG, and more realistic spatial modeling, possibly extended to whole-heart simulations including a surrounding torso like in [78, 79, 80, 81], could potentially improve the reliability of the computed QT intervals.

#### 4.3.2 Predicted adult drug responses for SQT1 patients

The estimated effects of the four drugs quinidine, ivabradine, ajmaline and mexiletine on the ventricular action potential and QT interval of adult SQT1 patients revealed that all four drugs are expected to lead to a significant increase in the action potential duration and the QT interval of SQT1 patients (see Figure 9). It is generally hard to validate the drug responses predicted by the computational procedure because data of drug responses for adult SQT1 patients would be required. However, for the drug quinidine, such experiments are reported in [15] in the form of measured QT intervals for a number of different heart rates. These measurements have been conducted for SQT1 patients without pharmacological treatment, SQT1 patients using the drug quinidine and for healthy adults without the SQT1 mutation [15]. We found that the effect of quinidine for adult patients estimated by the computational procedure fitted well with these measured drug responses (see Figure 10), indicating that the applied computational procedure shows promise for reliably predicting adult drug responses based on measurements of hiPSC-CMs.

### 4.4 Conclusions

In this paper, we have used a computational procedure to predict the effect of the drugs quinidine, ivabradine, ajmaline and mexilitine for clinical biomarkers of adult SQT1 patients, based on measured drug responses for hiPSC-CMs generated from an SQT1 patient. The procedure consists of identifying drug effects on individual ion channels based on membrane potential measurements of drug effects for hiPSC-CMs and then mapping this effect up to the adult case based on the assumption of functional invariance of ion channels during maturation. All four drugs were predicted to lead to significant increases in the ventricular action potential duration and QT interval. Furthermore, the effect of quinidine predicted by the computational procedure appeared to be in good agreement with measured effects of the drug. Consequently, we conclude that the computational procedure applied in this study could be a useful tool for estimating adult drug responses based on measurements of hiPSC-CMs, also as new measurements become available, including measurements of alternative drugs or mutations.

## Supporting Information

**S1 Appendix. Base model**. Base model formulation and singular value decomposition analysis of the base model currents.

